# Relative stability of different conformations and pathways of ligand escape in LIV-binding protein: A molecular dynamics simulation study

**DOI:** 10.1101/2024.10.20.619253

**Authors:** Jayita Das

## Abstract

Periplasmic binding proteins (PBPs) are a large family of receptors and transporters, present in gram-negative bacteria, which play a pivotal role in transport, chemotaxis and quorum sensing. These class of proteins have a distinct two domain architecture that undergo large conformational transitions through a hinge motion. In particular, Leucine-Isoleucine-Valine (LIV) binding protein can sense these specific side chains and undergoes a transition from a open to a closed conformation which is traditionally viewed as ligand induced conformational change. Here, atomistic molecular dynamics simulations and well-tempered metadynamics simulations are used to understand the role of ligand (namely, Isoleucine) in the conformational transition/selection of the LIV-binding protein. Furthermore, the pathway, energetics, and sequence of events during ligand escape/unbinding are unveiled from a microscopic perspective. Two distinct ligand binding pathways are identified: (i) domain separation followed by ligand escape, and (ii) ligand escape either followed by or synchronous with domain separation. The former is found to be energetically more favorable. Water is found to play an important role in the ligand unbinding process as well as providing stability to the closed conformation in the absence of the ligand, by forming bridging hydrogen bonds between two domains.

## Introduction

Bacteria sense small molecules and transport them into cytoplasm with the help of a receptor present in Periplasmic binding proteins (PBPs).^1–3^ These non enzymatic receptors are present in gram-negative bacteria (such as *Escherichia coli, Pseudomonas aeruginosa, Yersinia pestis* etc.) and exhibit dramatic conformational transitions. PBPs are found to be important in various cellular processes such as uptake and transport of many metabolically important solutes (amino acids, sugars and ions), chemotaxis and quorum sensing. They are also an integral component of ATP-binding cassette (ABC) transport systems. PBPs have a high affinity for their substrates, which is conferred by a set of hydrogen bonds that interact with the substrate. PBPs can also fulfill a biosensing niche for non-immunogenic targets.^1,4^

PBPs typically consist of two distinct domains connected by a hinge region. The ligand binding site is located in a cleft between the domains.^5,6^ Binding of the ligand induces a conformational change. This is commonly referred to as the “Venus flytrap” model of binding.^7^ This is characterized by an open conformation in the absence of the ligand and a closed conformation upon ligand binding. It is also possible that such PBPs exist as both open and closed forms in their ligand-free state and the ligand shifts the population towards the closed state. Such conformational change is crucial for their function. PBPs bind specific ligands with high affinity and specificity. Once the ligand is bound, the PBPs interacts with a membrane-associated ABC transporter. The energy for the transport process is provided by ATP hydrolysis, which facilitates the translocation of the ligand across the membrane.

The Leucine-Isoleucine-Valine (LIV) binding protein, known as the branched-chain amino acid binding protein, is a PBP involved in the transport of leucine, isoleucine, and valine, which are essential branched-chain amino acids. These amino acids serve as building blocks for proteins and as precursors for various metabolic pathways. The ability to import branched-chain amino acids is critical for bacterial growth and survival, especially in nutrient-limited environments.^8–11^ The structure of LIV-binding protein was elucidated using high-resolution X-ray crystallography, representing the protein in its superopen form (PDB ID: 1Z15) and ligand-bound forms (PDB IDs: 1Z16, 1Z17, and 1Z18).^5^ The binding site is located in a deep cleft between the two domains. The protein tightly encloses the amino acids, facilitating highly specific and high-affinity binding. The ligands are stabilized by a network of hydrogen bonds and hydrophobic interactions, ensuring precise recognition.^12^

Trakhanov *et al*. reported the ligand free and ligand bound structures of LIV-binding protein accompanied by normal mode analyses and short targeted molecular dynamics (TMD) simulations.^5^ They reported that the ‘closed’ structures with different ligands are very similar to each other whereas the ‘open’ (and superopen) form is vastly distinct from the close structure. The root mean squared deviation of the open and closed structures is approximately 6.5 Å.^13^ It is identified that the ligand is held together to the protein cleft with the help of several hydrogen bonds (HBs) and van der Waals (vdW) interactions. Residues Ser-79, Thr-102, and Ala-100 from domain-2 and Tyr-202 and Glu-226 from domain-1 form multiple Hydrogen bonding interactions with the ligands. Additionally, the ligand could get enthalpic stabilization through vdW interactions with residues Tyr-18, Leu-77, Leu-78, Tyr-150, and Phe-276.^5^ As the three ligand-bound structures are almost identical, the isoleucine bound structure is chosen in this study as a prototype of the closed form.

Studies involving conformational transitions of various proteins have been on the forefront of theoretical and experimental biophysics for several years.^14–19^ Although a detailed simulation study on LIV-binding protein is missing in the literature, similar PBPs were explored. Tang *et al*. used paramagnetic NMR on maltose binding protein (MBP) to discover that there exists a rapid exchange (in the ns to µs regime) between the open (∼ 95% population) and semi-closed (∼ 5% population) forms.^15^ McCammon and colleagues investigated MBP by using accelerated MD simulations and continuum electrostatics calculations.^14^ They reported the existance of a semi-closed conformation which is stabilized due to hydrophobic interactions. Consistent with earlier experimental results by Tang *et al*.,^15^ the open form is slightly more stable than the closed form, but with roughly same total energy. Based on the simulations they hypothesised a two step mechanism for the open → close state. The first step is the formation of a semi-closed state followed by the fast “ligand induced fit” mechanism. Yang and Mancera studied the lysine-arginine-ornithine (LAO) binding protein with elevated temperature MD simulations using CFF force-field.^20^ From the high-temperature simulations it was shown that the backbone peptide bond between Asp-11 and Thr-12 undergoes a large rotation during ornithin binding. Salopek-Sondi *et al*. investigated the role of residue number 18 in two PBPs namely, leucine-binding protein and LIV-bonding protein. Interestingly, they found the presence of Trp-18 disallows valine and isoleucine in case of leucine-binding protein.^21^ In LIV-binding protein, the same position is occupied by Tyr which allows all three amino acids.

Despite the earlier works, several important aspects has remained poorly understood, especially for LIV-binding protein. In this paper, the following questions are addressed: (i) Can the open conformation spontaneously convert into the closed form (and *vice versa*)? (ii) How important is the presence of ligand in the binding pocket to hold the closed form? In other words, can the closed form stabilize without the ligand? (iii) What is the free-energy barrier of ligand unbinding/binding? What is/are the pathway(s) and sequence of events of the same? In this paper, the above questions are answered with the help of both unbiased and biased atomistic MD simulations.

## Methods

The initial atomic coordinates were obtained from the protein data bank (PDB ID 1Z15 and 1Z17) for the open and ligand bound forms respectively.^5^ In this case the ligand is isoleucine (ILE). The required files for running MD simulation with GROMACS^22^ (version 2021.3) were obtained from CHARMM-GUI server.^23^ With these structures three different simulations were carried out: (i) The open form, (ii) the closed form but without the ligand, and (iii) the closed form with the ligand. In addition to these unbiased simulations five independent metadynamics simulations were run to estimate the free energy barrier of ligand escape and the sequence of events during the process.

All the proteins were placed inside a (10 nm)^3^ cubic box filled with TIP3P water and 0.15M Na^+^ and Cl^−^ ions. The box size is sufficient to avoid the periodic image interaction. Periodic boundary conditions were imposed in all three directions. The systems were first energy minimized followed by 100 ps equilibration by position restraining the protein. Then a 100 ns production run were performed. All the simulations used the CHARMM36m force field^24^ in NpT ensemble (p=1 bar and T=300 K). To fix the temperature around 300 K modified Berendsen thermostat was used (*τ*_*T*_ =1.0 ps) and to keep the average pressure around 1 bar C-rescale barostat was used (*τ*_*P*_ =5.0 ps and compressibility 4.5 *×* 10^−5^ bar^−1^). The leap-frog algorithm was used to integrate the Newton’s equations of motion with a timestep of 1 fs (for equilibration) and 2 fs (for production run). The coordinates were dumped in every 10 ps. The Verlet cutoff scheme was used. For vdW and short range electrostatic interaction a 1.2 nm cutoff was used. For long range electrostatic interaction Particle Mesh Ewald (PME) algorithm was employed with an FFT grid spacing of 0.12 nm. All bonded interactions were constrained using the LINCS algorithm.

To find out the free-energy barrier of the ligand (here, Ile) escape from the binding site, well-tempered metadynamics simulations^25^ were performed using PLUMED^26^ (patched with GROMACS). The collective variable was defined to be the distance between the *C*_*α*_ of Glu-328 and the center-of-mass (COM) of the ligand. The *C*_*α*_ of Glu-328 resides at the junction of the two domains and acts as a fulcrum to the hinge motion (red bead in Figure 1). Five independent metadynamics run were performed to check the robustness. The hill deposition frequency was set to 500 steps with a bias-factor of 5. An upper wall was placed at d=5 nm, to prevent sampling of uninteresting regions. All other settings were the same as described before. The metadynamics simulations were truncated upto the time when the ligand leaves the binding site. Of course, this cannot give us a converged free-energy landscape. However, it can be useful to extract the free-energy barrier of ligand escape.

**Figure 1:**
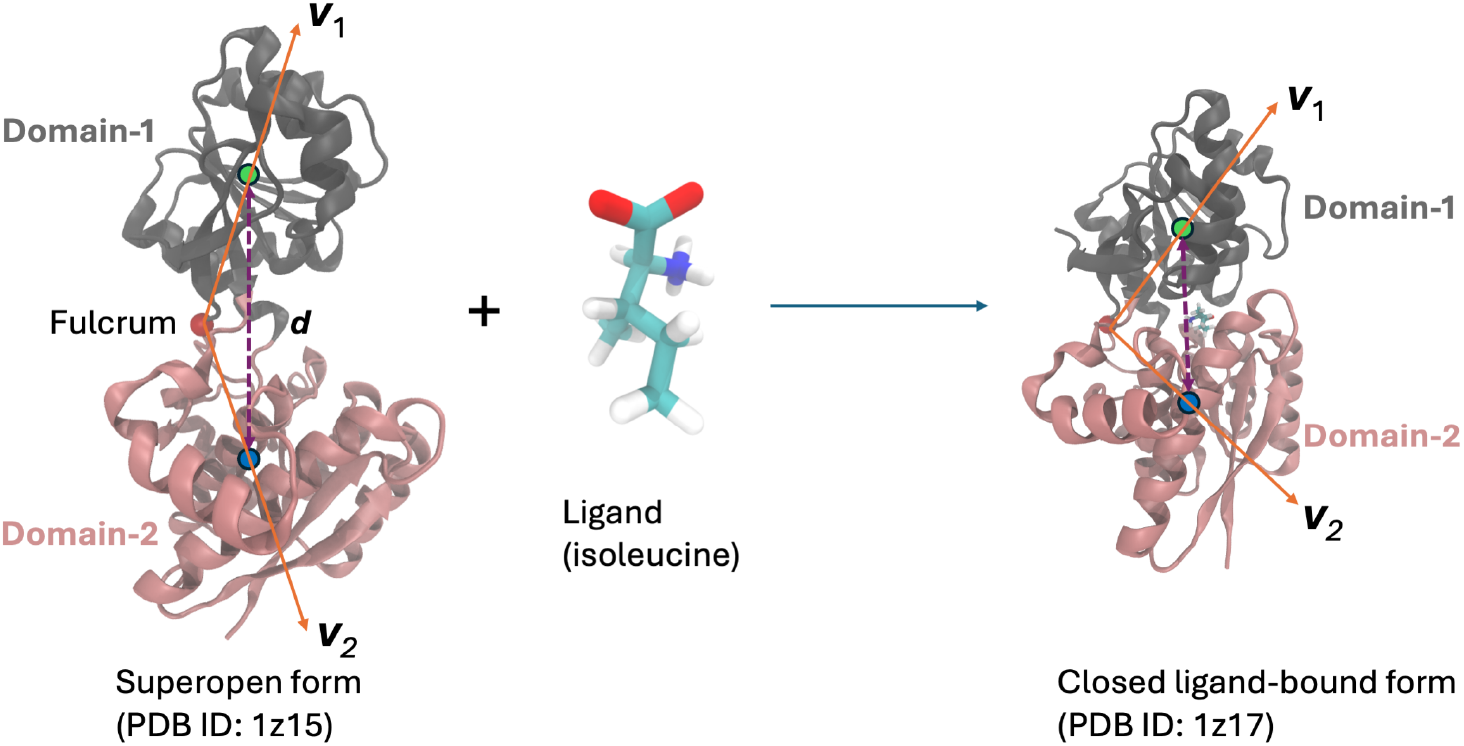
The transition between open/superopen (PDB ID: 1z15) and closed/ligand-bound (PDB ID: 1z17) forms.^5^ The protein structure can be divided into two distinct lobes shown in grey (domain-1) and pink (domain-2). The lobes show hinge movement around a fulcrum (red sphere). The distance between the centers of mass (COMs) as well as the angle between the two vectors (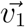 and 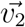) can be used as order parameters to quantify open and closed conformations. To study the ligand binding/unbinding, the distance of the ligand from the fulcrum is used.

The analyses were done using the in-built tools in GROMACS and sometimes using own Python scripts. The visualization was done using Visual Molecular Dynamics (VMD)^27^ software and graphs were created using Python.^28^

## Results and discussion

### Structure and dynamics of open and closed conformations

LIV-binding protein adopts a bi-lobed arrangement connected by a flexible hinge region. The two globular domains, referred to as domain-1 (resiues 121-248 and 329-344) and domain-2 (residues 1-120 and 249-328) in this study [Figure 1], form a deep binding cleft where the ligand is accommodated. Upon ligand binding, the protein undergoes a significant conformational change, transitioning from an open to a closed state. From the X-ray crystallography structures, the inter-domain center of mass (COM) distance for the superopen form is ∼ 3.4 nm and that of the ligand-bound closed form is ∼ 2.77 nm. In the absence of a ligand, LIV-binding protein predominantly exists in an open conformation, with the two lobes positioned apart, allowing for rapid scanning and binding of branched-chain amino acids.

From the 100 ns unbiased MD simulation trajectories, root mean squared deviations (RMSD) of each domain as well as the full protein are calculated [Figure 2]. Domain-1 of the super-open form is found to be more dynamic than those of closed (with ligand) and closed (without ligand) conformations [Figure 2(a)]. But, RMSD of domain-2 initially showed similar mobility for all the three systems [Figure 2(b)]. After 70 ns simulation time, RMSD of domain-2 of the closed (without ligand) system shows a marked increse. In Figure 2(c), the RMSD of the whole proteins are plotted. It is observed that the RMSD of the super-open form periodically varies with time. The closed (with ligand) protein showed almost no change during the simulation and RMSD of the closed (without ligand) shows an increase above 70 ns.

**Figure 2:**
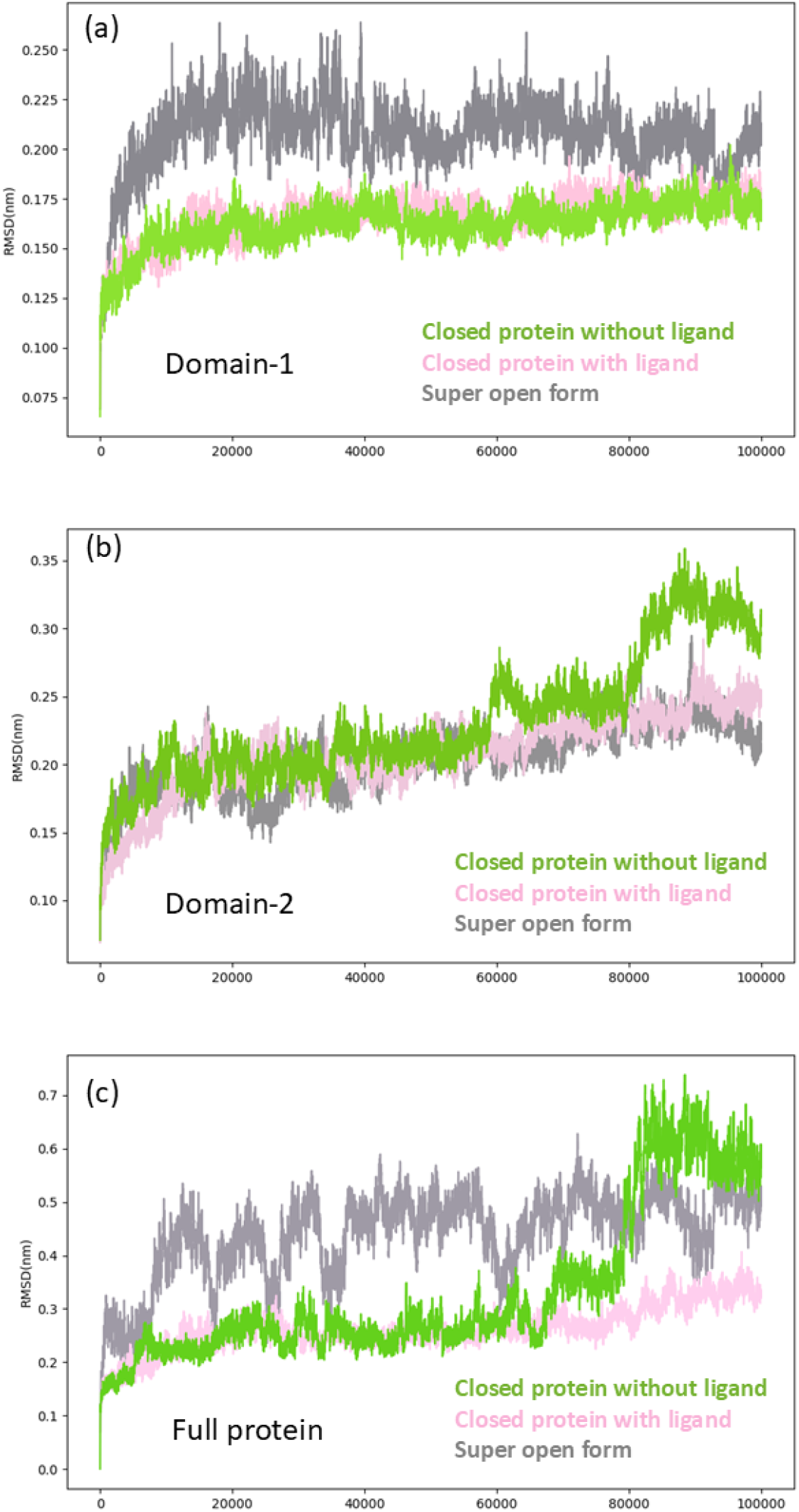
Root Mean Squared Deviation (RMSD) of (a) domain-1, (b) domain-2, and (c) the whole protein for 1z15 super open form (gray), 1z17 closed form with ligand (light pink) and 1z17 closed form without ligand (yellow-green).

To check for the hinge motion, the inter-domain COM separation distance and the angle between two domains with respect to the fulcrum are calculated against simulation time and plotted in Figure 3(a) and Figure 3(b) respectively. The inter-domain COM distance for the super-open form starts around 3.4 nm and eventually decreases to near 3.0 nm, indicating an open form. The closed (with ligand) form maintains its inter-domain distance around 2.85 nm which is very close to the X-ray crystallography value of 2.77 nm. The closed (without ligand) conformation maintained its close form up to 70 ns and then sharply increases to 3.2 nm indicating an open form. The inter-domain angle variation tells the same story: the superopen remains open with more fluctiuations, the closed (with ligand) form maintains the inter-domain angle between 90 and 100 degrees, and the closed (without ligand) form shows a sharp increase in the inter-domain angle beyond 70 ns of the simulation time.

**Figure 3:**
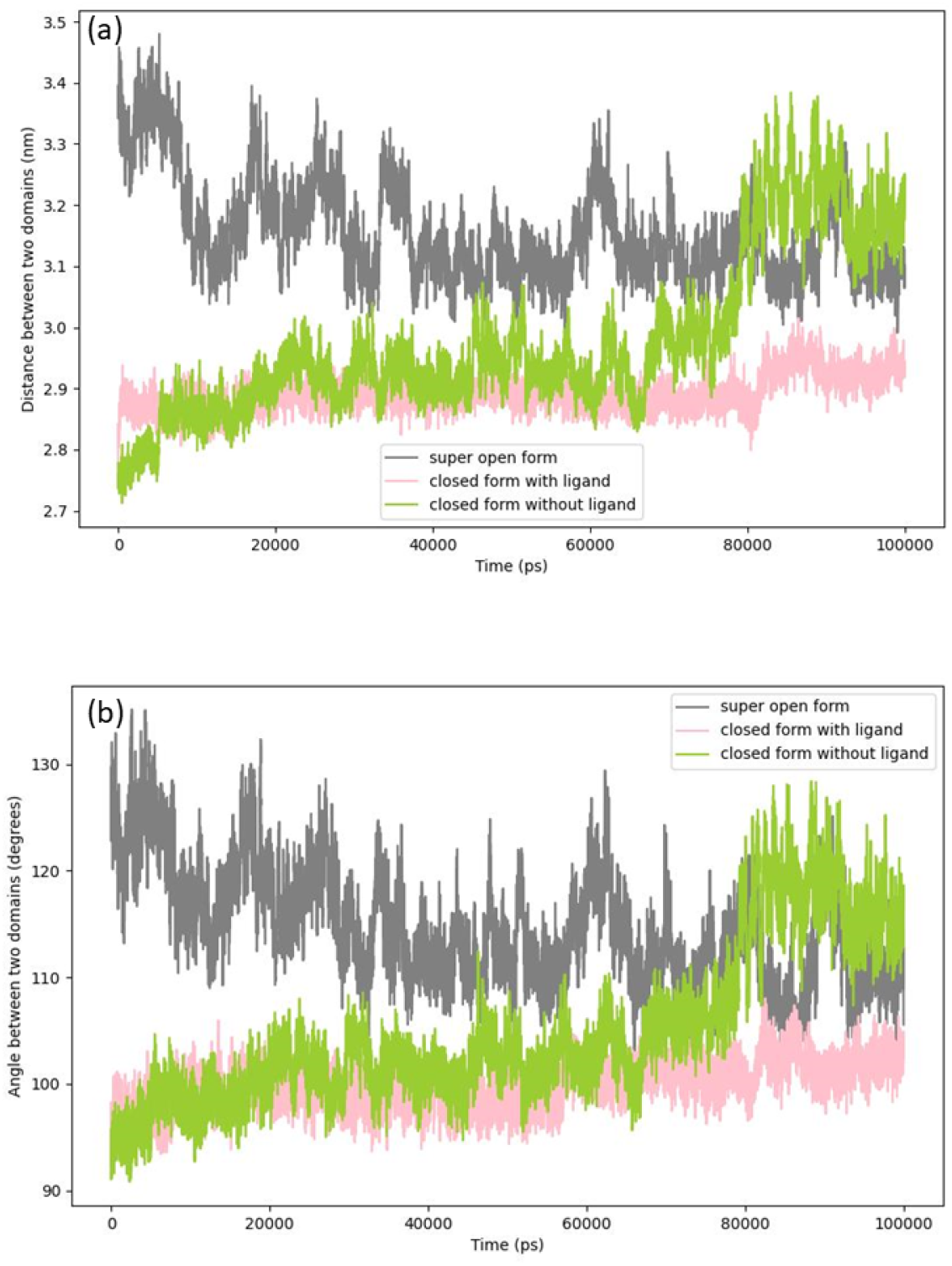
(a) Inter-domain separation distance [shown in Figure 1] for open, closed (with ligand), and closed (without ligand) along the 100 ns unbiased simulation trajectory. (b) Inter-domain angle between two vectors connecting the fulcrum with the COMs of each domain as again described in Figure 1.

The ligand (Ile) is held at the cleft region between two domains by non-bonded interactions which include electrostatic and hydrophobic attractions. The former appears in the form of hydrogen bonding (HB) which plays an important role in providing the much needed enthalpic stabilization. As discussed in the introduction, Ser-79, Thr-102, and Ala-100, Tyr-202 and Glu-226 form multiple HB with the ligands. In a way, the ligand acts as an anchor that keeps the two domains together. The general concensus is that the ligand shifts the conformational ensemble to the closed from and without ligand the protein stays as the open/super-open/semi-open form.^14,15^ However, from Figure 3 it can be seen that the closed form can survive for several tens of nanoseconds without the ligand acting as an anchor between the two domains. More insight on the origin of its stability will be discussed later.

### Ligand escape pathway, energetics, and sequence of events

The ligand bound state is stable and the ligand seem to be tightly bound through non-bonded interactions as discussed before. Hence, it is almost impossible to see a ligand escaping event through the unbiased simulations, making it a rare event.^29^ With the help of metadynamics, one can study the energetics of ligand escape, the possible pathways, and sequence of events during the escape. Here we bias one (slower) collective variable (distance between Glu-328 *C*_*α*_ and COM of the ligand) as discussed in the methods section and kept the (faster) collective variables (such as water coordination, domain movement etc.) unbiased.

Based on the five independent metadynamics trajectories, two distinct escape pathways/mechanisms were observed. In one pathway (path-I), as graphically shown in Figure 4(a), 4(b), and 4(c), the domains first move away from each other exposing the ligand as well as the cleft region to the solvent [Figure 4(b)]. In this state, the ligand remained bound to the domain-2, hydrogen bonded with Ser-79 and Thr-102. Here the ligand got partially solvated by water that entered due to the domain movement. Later, the ligand detaches from domain-2 [Figure 4(c)] and becomes fully solvated.

**Figure 4:**
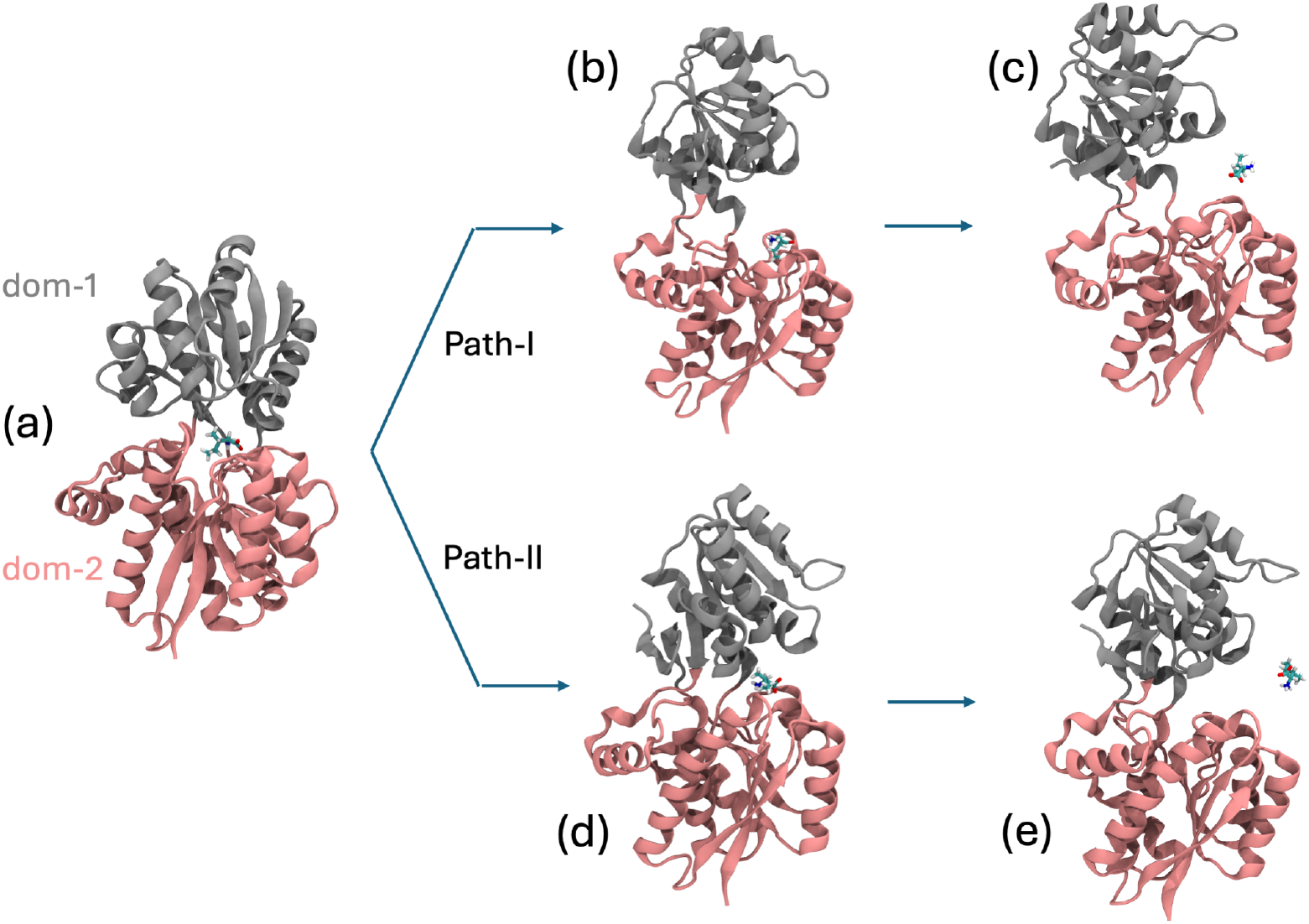
Pathway-I and II of ligand (isoleucine) escape from the cleft of LIV-binding protein. The whole protein is devided into domain-1 (in grey) and domain-2 (in pink). (a) Initial ligand bound state, (b) Intermediate of path-I: Domain separation where the ligand is attached to domain-2, (c) ligand excape and relaxation of protein conformation, (d) Intermediate of path-II: Domains are partially opened and ligand is in contact with both the domains. (e) ligand excape and relaxation of protein conformation for path-II.

In Figures 5(a)-5(c), path-I is shown with the help of two variables: (i) the distances between the ligand and the fulcrum of the protein (*C*_*α*_ of Glu-328) which is also the collective variable for metadynamics along which the system is biased, and (ii) the inter-domain separation distance which is unbiased. It can be seen that in all three cases, the domain separation occurs before the ligand escape.

**Figure 5:**
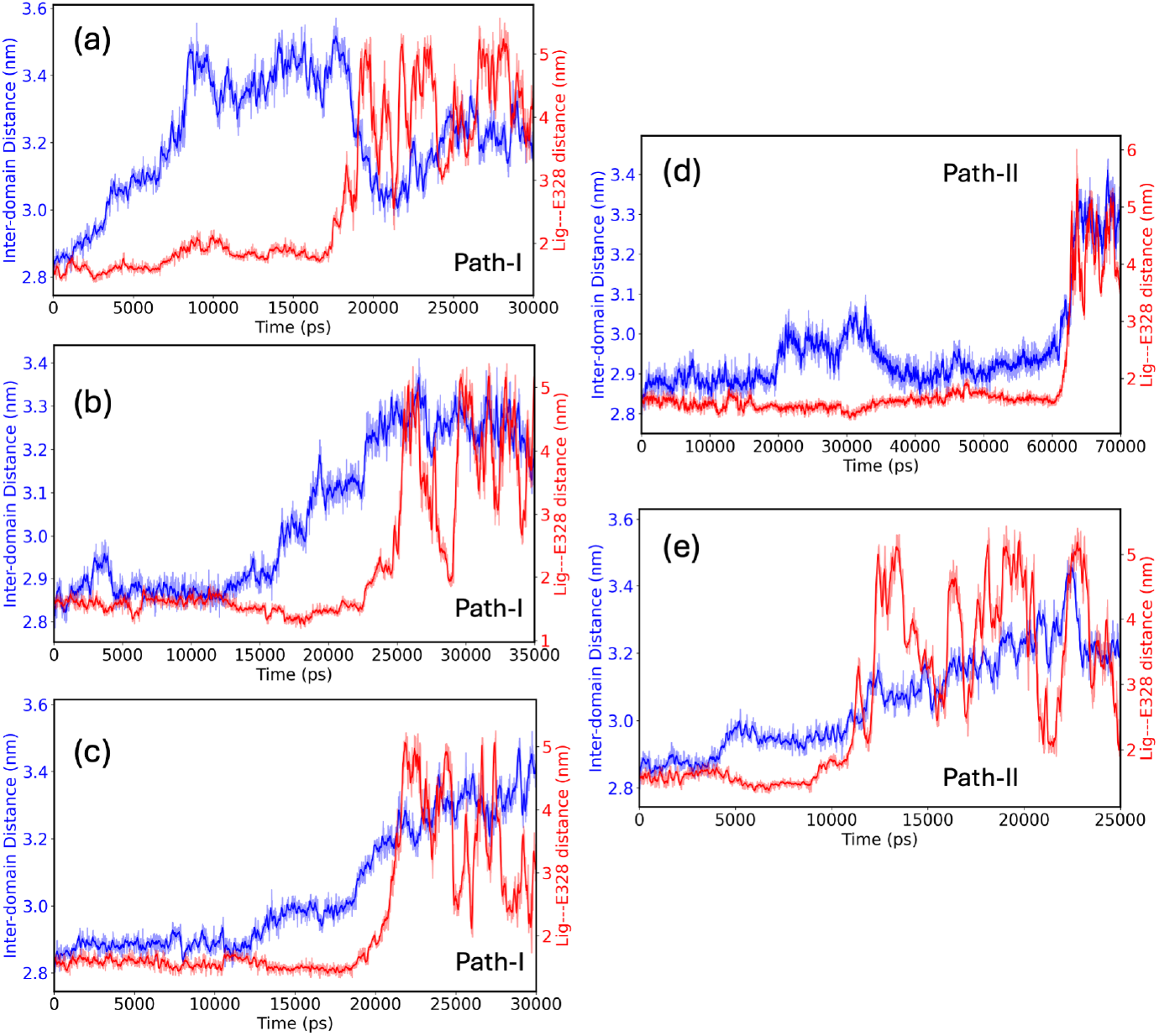
Demonstration of two different ligand escape pathways. The inter-domain COM distances are plotted in blue and the distances between the protein fulcrum and the ligand are plotted in red. Panels (a), (b), and (c) shows that domain separation occurs before the ligand has escaped from the cleft. On the other hand, panels (d) and (e) shows that either ligand escape and domain separation occur synchronously or ligand escape happens before the domain separation.

In another pathway (path-II), as shown in Figure 4(a), 4(d), and 4(e) the ligand escape either occurs without the domain opening [Figure 5(e)] or it happens synchronously [Figure 5(d)]. From the metadynamics simulations up to the transition point, path-I (Δ*G*^‡^ ≃ 40*kJ/mol*) is found to be energetically more favourable than path-II (Δ*G*^‡^ ≃ 60*kJ/mol*).

In a previous targeted-molecular-dynamics study, Trakhanov *et al*. studied ligand unbinding from LIV-binding protein and found the domain-2 bound state (i.e., Ile H-bonded to S79 and T102) before the unbinding happened. ^5^ However, they missed the other pathway and also could not comment on the free-energy barrier of ligand escape.

### Role of water

Water plays an important role in biological processes.^30–33^ Water acts as a lubricant that facilitates biochemical reactions and stabilizes macromolecular structures through hydrogen bonding. Its ability to form dynamic hydration shells around biomolecules, such as proteins and nucleic acids, is essential for maintaining their proper folding, function, and interactions. Additionally, water’s unique properties enable the efficient transport of ions and molecules, making it fundamental to cellular processes and overall life.

This section is devoted to understand the role of molecular water in the conformational fluctuation of LIV-binding protein and also the ligand escape pathway. In the context of another conformation shifting protein, Adenylate kinase, an earlier study by Adkar *et al*. revealed the presence of a half-open/half-closed (HOHC) state, produced by the relative movement of the LID and NMP domains.^16^ The stability of the HOHC state was attributed to the increased interaction between polar amino acid side chains with water. This reduces the conformational free energy barrier by ∼ 20 kJ/mol. Interestingly enough, in the present system of interest, the ligand unbinding free-energy barrier also gets reduced by ∼ 20 kJ/mol when the intermediate state goes through a semi-open form [Figure 4(b)].

In the ligand escape metadynamics simulations two distinct pathways were observed that have been described in the previous section. In both of them water molecules were found to stabilize the intermediate state before the unbinding takes places. In pathway-I the intermediate is the open form where the ligand is attached only to domain-2. In that state water molecules partially solvate the ligand [Figure 6(b)] as well as the polar side chain of domain-1. Subsequently when the ligand fully escapes the pocket water molecules completely solvate the ligand by forming several hydrogen bonds [Figure 6(c)]. In the second pathway the intermediate is not a fully open conformation therefore the ligand retains majority of its hydrogen bonding interactions with both the domains. However here also a few water molecules form stable hydrogen bonding interaction with the ligand and also bridging bonds as shown in Figure 6(d). Similar to pathway-I the ligand escape is supported by a complete solvation of the ligand [Figure 6(e)]. The ligand unbinding, unlike any other process, relies on the entropy-enthalpy interplay. Although it may look like the driving force is enthalpic stabilization of the ligand as well as the amino-acid sidechains at the cleft by water, a decomposition of the free energy into enthalpic (Δ*H*) and entropic (*T*Δ*S*) components are necessary.^34^

**Figure 6:**
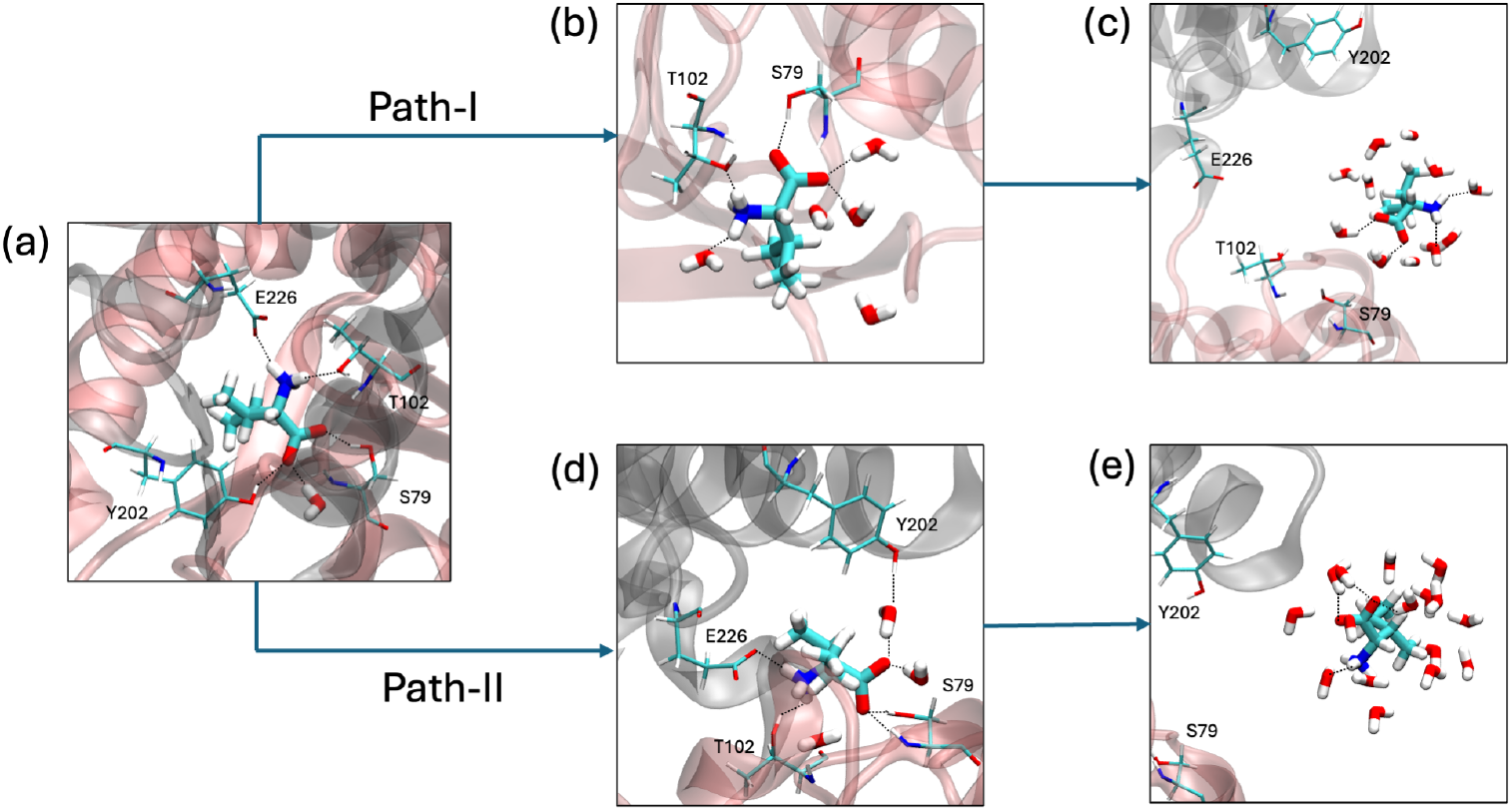
Magnified view of the ligand and its surroundings for the two ligand escape pathways. (a) The initial state where the ligand (isoleucine) is H-bonded to four amino acid sidechains: S79, T102, Y202, and E226. (b) The intermediate of path-I where the ligand is attached to domain-2 and H-bonded with S79, T102, and partially solvated by a few water molecules. (c) Full escape of the ligand solvated by water molecules. (d) The intermediate of path-II where the ligand retained H-bonded interaction with three out of the four sidechains: S79, T102, and E226. On top of that a water mediated bridging H-bond is formed between the ligand and Y202. (e) same as ‘c’ for path-II.

From the unbiased simulations, important information regarding the role of water can be extracted. In Figure 7(a), The radial distribution functions (RDFs) between one of the cleft amino acids (Ser-79) and the oxygen atom of water are plotted. From the RDFs one can see that the closed form but without ligand has a significant amount of water molecules in the cleft region. On the other hand the cleft region is almost dry when the ligand is present. It is understandable that the ligand favourably interacts with both the domains and tries to keep them close to one another. In this way the ligand bound closed form gains its stability. This aspect has been well established in previous studies for similar systems.^14^

**Figure 7:**
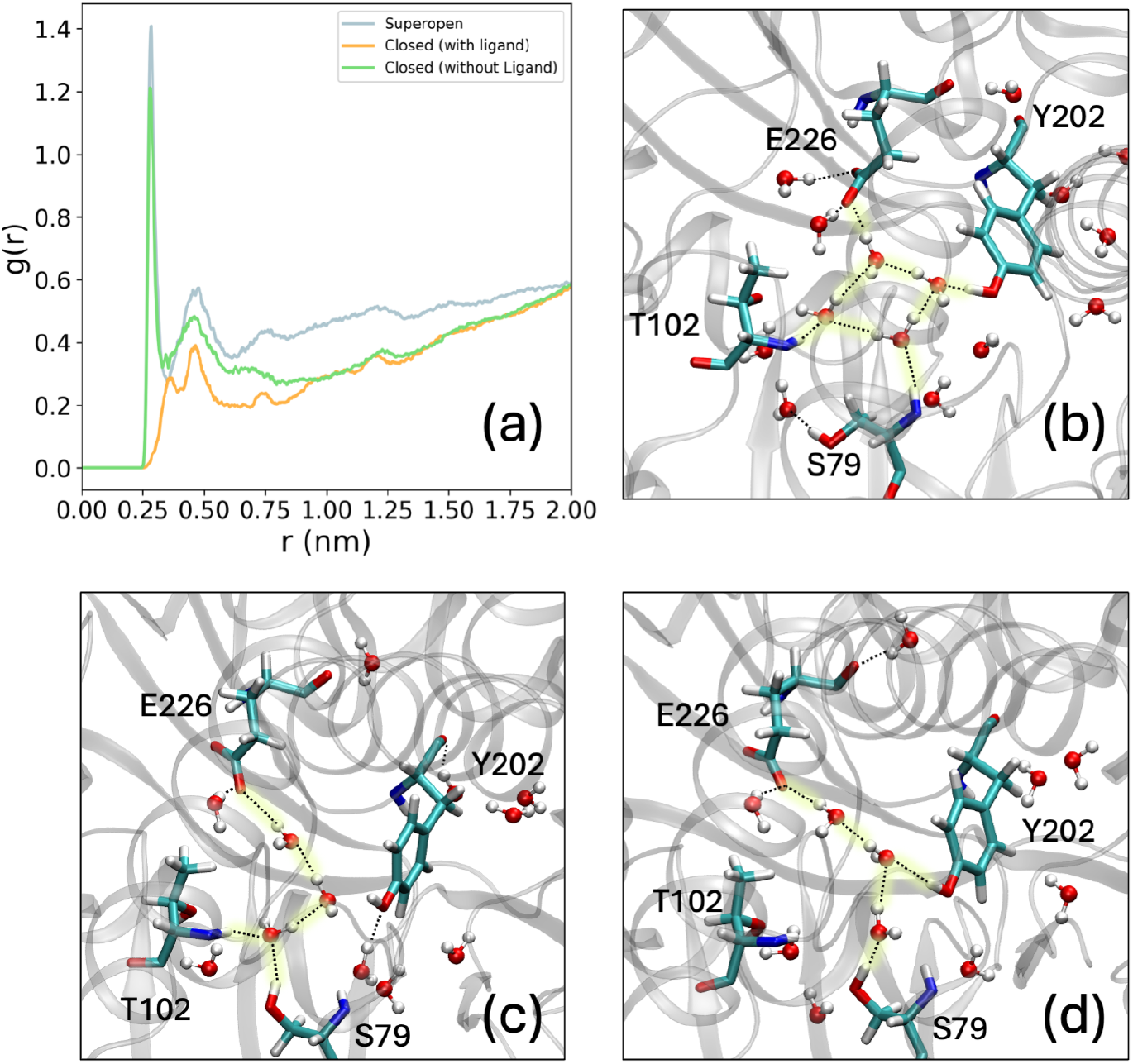
(a) Radial distribution function between S79 and water from unbiased trajectories for the superopen, closed (with ligand), and closed (without ligand) conformations. (b), (c) and (d) are representation of water mediated bridge hydrogen bonding (highlighted in yellow) between E226, Y202, S79, and T102 in the closed form of the protein without ligand. Such interaction acts as anchors between the two domains of the protein and makes the closed state stable in the absence of the ligand. The H-bonding interaction was observed by using the geometric criteria given by Luzar *et al*. with *r*_*cut*_ = 3.0*Å* and *θ*_*cut*_ = 30 *degrees*.^35^

However the unanswered question is: *how the closed conformation remain closed even if there is no ligand in between the domains*. It has been found that several water molecules infiltrate the cleft region and form extended water mediated bridging hydrogen bonds as shown by some representative snapshots in Figure 7(b), 7(c) and7(d). In Figure 7(b), water molecules can be seen to form a four membered ring whose four vertices are H-bonded to four different amino acid sidechains. Therefore, water molecules play an active role in the stability of the closed confirmation without the presence of a the ligand. The importance of water mediated interactions at protein-protein interface was explored before.^36,37^

## Conclusion

The conformational fluctuations of LIV binding protein and its interaction with ligands were investigated, with a focus on the dynamics of ligand escape. Previous assumptions suggested that the ligand was necessary to sustain the closed conformation of LIV binding protein. However, the results of this study demonstrate that the closed conformation can remain stable without the ligand, primarily due to the stabilizing effect of water-mediated hydrogen bonds bridging key sidechains of domain-1 and domain-2. This finding challenges the conventional view and provides new insights into the structural behavior of the protein under ligand-free conditions.

Two distinct pathways for ligand escape were identified. In one pathway, the ligand escapes after domain opening, while in the other, ligand escape occurs either simultaneously with or prior to domain opening. It was also observed that the pathway where domain opening precedes ligand escape has a lower free energy barrier, highlighting a more energetically favorable route for ligand release.

Looking forward, further studies could explore the effects of other ligands, such as valine and leucine, on the conformational dynamics and ligand escape pathways of the protein. Additionally, a deeper investigation into the entropy-enthalpy balance and kinetics of these processes could provide a more comprehensive understanding of the underlying mechanisms. Such insights may offer valuable information for the development of targeted inhibitors or modulators of ligand binding and release.

## Acknowledgement

I would like to express my sincere gratitude to the SMCB seminar series (https://sites.google.com/view/smcb2021) for inspiring me to explore theoretical and computational biophysics in greater depth, which ultimately motivated the inception of this work. I am also deeply thankful to the INSPIRE fellowship (DST, India) for their financial support at the start of this project. I am grateful to Prof. John Straub and Prof. Qiang Cui, whose courses at Boston University significantly deepened my understanding of statistical mechanics, coding, and data analysis, all of which were invaluable to this research. Lastly, I extend my thanks to Dr. Sayantan Mondal for his guidance in manuscript preparation and for the valuable discussions.

